# m6A RNA methylation attenuates thermotolerance in Arabidopsis

**DOI:** 10.1101/2025.05.22.655480

**Authors:** Kirti Shekhawat, Arsheed H. Sheikh, Kashif Nawaz, Anam Fatima, Waad Alzayed, Arun Prasanna Nagarajan, Heribert Hirt

## Abstract

Heat stress severely affects plant growth, causing significant crop yield losses. To counteract this, plants employ adaptive strategies, including chromatin and transcriptional regulation of heat stress-related genes. However, the role of RNA methylation in heat stress response remains unclear. This study identifies m6A RNA methylation as a negative regulator of thermotolerance in Arabidopsis. Loss of m6A modification enhances resilience to heat stress by increasing transcript levels of heat-responsive genes and stabilizing their mRNAs. Upon heat shock, plants transiently reduce m6A levels by modulating key methylation machinery genes. This m6A deficiency enriches the H3K4me3 histone mark at heat-responsive gene loci, boosting their transcription. The combined effect of m6A-regulated chromatin changes and mRNA stability facilitates the accumulation of heat shock proteins essential for cellular protection and stress response. These findings reveal an interplay between m6A RNA methylation and histone modifications, forming a regulatory network that fine-tunes gene expression during heat stress. This novel mechanism highlights m6A as a critical regulator of plant thermotolerance, offering a promising target for improving heat stress resilience in crops.

## Introduction

N6-methyladenosine (m6A) is the most abundant internal modification of mRNA in eukaryotes, and it has emerged as a critical regulatory element in modulating various physiological processes, including responses to abiotic stresses. This modification is dynamically regulated by a set of proteins known as “writers” (methyltransferases), “erasers” (demethylases), and “readers” (m6A-binding proteins), which add, remove, or recognize the m6A mark, respectively (Arribas-Hernández *et al*., 2018; Reichel, Köster and Staiger, 2019). While m6A modification has been extensively studied for its role in controlling plant growth and development, recent studies have specifically highlighted its importance in conferring drought, salt tolerance and dark induced senescence in plants, suggesting that m6A may act as a pivotal regulator of abiotic stress resilience (Ganguly *et al*., 2024; Hou *et al*., 2022; Hu *et al*., 2021; Sheikh *et al*., 2024).

Despite these advances, the role of m6A modification in plant thermotolerance remains largely unexplored. Thermotolerance is an essential adaptive trait for plants, especially given the rising global temperatures and frequent heat waves that pose a significant threat to crop productivity and yield. Upon exposure to elevated temperatures, plants rapidly activate heat shock transcription factors (*HSFs*), which subsequently upregulate the expression of heat shock proteins (*HSPs*) (Bakery *et al*., 2024; Huang *et al*., 2023; Lämke *et al*., 2016; Shekhawat *et al*., 2021). These HSPs act as molecular chaperones that stabilize and refold denatured proteins, protecting the plant from heat-induced damage and ensuring cellular homeostasis under stressful conditions (Kotak *et al*., 2007; Wang *et al*., 2023; Yoshida *et al*., 2011). The controlled regulation of HSPs is important for the efficient recovery from the heat-shock in plants. Beyond the immediate response to heat stress, plants also rely on thermomemory, allowing them to “remember” prior heat stress events and respond more efficiently to subsequent exposures. This acquired thermotolerance is regulated through epigenetic mechanisms, with histone modifications such as H3K4me2 and H3K4me3 playing a key role in establishing and maintaining the memory of heat stress at specific gene loci, often referred to as heat stress memory genes. The transcription factors HSFA2 and HSFA3 are central to this process, as they facilitate the recruitment of H3K4me3 modifications to these loci, thereby priming the genes for rapid reactivation upon subsequent heat stress encounters (Balazadeh, 2022; Friedrich *et al*., 2021; Kappel *et al*., 2023; Lämke, Brzezinka and Bäurle, 2016; Liu *et al*., 2019; Nishio, Kawakatsu and Yamaguchi, 2024; Shekhawat *et al*., 2022).

Given that m6As is involved in chromatin regulation in mammalian systems and also regulates the stability of stress responsive transcripts (Chen *et al*., 2024; Liu *et al*., 2020; Yu *et al*., 2021), we investigated whether m6A modification plays a role in heat stress response in plants. We observed higher resistance of m6A-deficient mutants to heat stress and that m6A levels are dynamically regulated during the heat shock response in Arabidopsis. The increased stability of *HSP* transcripts, along with their primed H3K4me3 chromatin states, contributes to enhanced thermotolerance revealing that m6A acts as a negative regulator of thermotolerance. This knowledge can give important leads for developing crops that can better withstand rising temperatures, which is crucial for maintaining agricultural productivity in the face of climate change.

## Material and Methods

### Plant Material Heat stress experiments

Arabidopsis (Arabidopsis thaliana) ecotype Col-0 and mta ABI3:MTA (*mta*) were used in this study. *mta* mutant was generated as described earlier (Bodi *et al*., 2012). MTA complementation line was generated by Agrobacterium (Agrobacterium tumefaciens) mediated transformation of *mta* plants by MTA genomic locus cloned into pGWB440 vector. Seeds were surface-sterilized for 10 min with 0.05% SDS solution prepared in 70% ethanol, followed by three times washing with absolute ethanol. The sterilized seeds were plated on ½ MS medium agar plates. Seeds were stratified at 4°C for 2 days and then plates were transferred to a growth chambers (Model CU36-L5, Percival Scientific, Perry, IA, USA) under a 16-h photoperiod and 8-h dark conditions at 22°C for germination and seedling growth. We gave a heat treatment by modifying a previous method (Larkindale and Vierling, 2008). 5-day-old seedlings of near equal lengths were transferred to new ½ MS plates. For HS treatments, 11 days old plants were exposed directly to 44°C HS for 30 min, for this 44°C HS treatment, we used a pre-heated water bath. For control treatment, plants were grown under normal conditions at 22°C (NHS).

### Measurement of chlorophyll content

Chlorophylls A and B were extracted from frozen shoot samples with 80% acetone(Arnon, 1949; Sims and Gamon, 2002). After centrifugation, the samples were centrifuged, and the chlorophyll concentration of supernatant was calculated using a spectrophotometer (Tecan). The supernatant was used to measure OD at 537, 647, 645, and 663 nm. The following formulas were used to calculate chlorophyll A/B: chlorophyll A (μmol ml−1) = 0.01373 A663 − 0.000897 A537 − 0.003046 A647; chlorophyll B (μmol ml−1) = 0.02405 A647 − 0.004305 A537 − 0.005507 A663 (Shekhawat *et al*., 2024).

### Ion leakage assay

The HS treated and NHS treated plants were incubated in 2.5 mM MES buffer containing 0.05% (v/v) Tween-20. The samples were incubated at 30°C in dark for 24 hours and readings were taken using a Conductivity meter (Seven Excellence, Mettler Toledo)(Sheikh *et al*., 2023).

### Total RNA and mRNA extraction from seedlings

Total RNA was extracted from the Arabidopsis seedlings grown on ½ MS with the Nucleospin RNA plant kit (Macherey-Nagel#740949) following the manufacturer’s recommendations. The quality and quantity of the RNA were assessed using Nanodrop-6000 spectrophotometer, 2100-Bioanalyzer (RNA integrity number greater than 8.0). poly(A+) + mRNA was isolated from 100 µg of total RNA using Oligo-dT dynabeads from mRNA isolation kit (Thermo#61006).

### RNA sequencing

We performed RNA sequencing on 4-wk-old adult Col-0 and *mta* plants was with three biological replicates. Briefly, RNA from HS and NHS treated plants was extracted using Nucleospin RNA plant kit (Macherey-Nagel) following the manufacturer’s recommendations. The quality and quantity of the RNA was assessed using Nanodrop-6000 spectrophotometer, 2100-Bioanalyzer (RNA integrity number greater than 8.0). RNA-seq was conducted using the Illumina TruSeq Standard mRNA Library Preparation protocol, following the manufacturer’s instructions for 150 bp paired-end sequencing. Using the Illumina HiSeq 4000 platform, pooled libraries were sequenced, followed by read trimming and alignment. Differential expression analysis was performed using DESeq2 with an FDR ≤ 0.01 (Love, Huber, and Anders, 2014), and functional enrichment of the identified DEGs was carried out via AgriGO using standard parameters (Tian *et al*., 2017).

### Dot blot assay

To quantify the relative levels of m^6^A, poly(A^+^)-enriched mRNA samples (200, 100, and 50 ng) were spotted onto SensiBlot Plus Nylon membranes (Fermentas #M1002) and air-dried for 5 minutes. The membranes were UV crosslinked using a Stratalinker and blocked for 3 hours in PBST containing 5% (w/v) non-fat milk. Following blocking, membranes were incubated overnight at 4°C with an anti-m^6^A antibody (Abcam #15320, 1:2500 dilution). After washing, membranes were treated with HRP-conjugated goat anti-rabbit secondary antibody (Promega, 1:10,000 dilution). Chemiluminescent signals were developed using the Immobilon Femto Western HRP substrate (Thermo #34094).

### Western blot

Two-week-old WT and *mta* mutant plants were harvested and immediately flash-frozen in liquid nitrogen. Nuclei extracts were mixed with 5× SDS loading dye, boiled at 85 °C for 10 minutes, and loaded onto a 10% (w/v) SDS-PAGE gel. Proteins were transferred to a PVDF membrane, which was blocked with TBST containing 5% (w/v) milk for 2 hours. The blot was then incubated overnight with a primary antibody (1:2500; Abcam) in TBST with 5% (w/v) milk. Following five washes, the membrane was incubated with an anti-rabbit secondary antibody (1:10,000; Promega) for 1 hour. After another five washes, the membrane was developed using ECL Clarity Max solution (Bio-Rad).

### Gene expression analysis by reverse transcription quantitative PCR

Total RNA was extracted and reverse transcribed into cDNA using the SuperScript III First-Strand Synthesis SuperMix kit (Invitrogen), following the manufacturer’s instructions. Reverse transcription quantitative PCR (RT-qPCR) was conducted on a CFX384 Real-Time PCR Detection System (Bio-Rad) using fivefold diluted cDNA as a template. Reactions were performed with Universal SYBR Green Supermix (Bio-Rad) under the following thermal conditions: 50⍰°C for 2⍰min, 95⍰°C for 10⍰min, followed by 40 cycles of 95⍰°C for 10⍰s and 60⍰°C for 40⍰s. A melt curve analysis was included to confirm the specificity of amplification. Data were analyzed using Bio-Rad CFX Manager software. All reactions were performed in technical triplicates, and Ct values were normalized to the internal reference gene Ubiquitin.

### Transcript stability time course

To assess mRNA stability, 11-day-old seedlings were transferred to ½ MS liquid medium and subjected to heat stress at 44⍰°C for 30 minutes. After a 24-hour recovery period, Actinomycin D (30⍰μg/mL; Sigma) was added to inhibit transcription. Samples were collected at 0, 3, and 6 hours post-treatment and immediately flash-frozen in liquid nitrogen (S Fig. 3A). Total RNA was isolated, and RT-qPCR was performed as described above. Transcript levels at 0 hours were set to 100%, and relative RNA abundance was calculated for 3- and 6-hour time points.

### Chromatin Immunoprecipitation (ChIP)

Chromatin immunoprecipitation (ChIP) was performed as previously described by Lämke *et al*. (17), with minor modifications. Briefly, 500⍰mg of 11-day-old Arabidopsis seedlings were cross-linked with 1% formaldehyde, quenched using glycine, and subsequently frozen at −80⍰°C. Nuclei were isolated and chromatin was sheared to approximately 250⍰bp fragments via sonication. Immunoprecipitation was carried out overnight at 4⍰°C using antibodies against H3 (ab1791) or H3K4me3 (ab8580), both pre-bound to protein A agarose beads. After extensive washing and reversal of cross-links, DNA was purified via ethanol precipitation and resuspended in nuclease-free water. ChIP-enrichment was quantified by qPCR, and values were normalized to total H3 levels. Data represent averages from three independent biological replicates.

## Results

### The m6A machinery is affected during heat stress

To investigate the impact of heat stress on plant m6A methylation, we analyzed global m6A levels under heat stress conditions using a dot blot assay. We exposed 11 days old Col-0 seedlings to a heat stress of 44°C for 30 minutes and harvested samples after 1h post heat stress. We observed a significant decrease in global m6A levels at 1 h heat-stressed plants compared to non-heat-stressed plants (Fig. 1A, B). This decrease in m6A levels at 1h correlated with a decrease in the transcript levels of prominent m6A readers and writers as reflected by qRT-PCR. We observed a significant decrease in transcript levels of *MTA, MTB, HAKAI*, and *VIR* the primary components of the m6A writer complex, after 1 hour of heat stress (Fig. 1C, S Fig. 1). Interestingly, these levels were restored at 24 hours post HS. Similarly, the transcript levels of the reader complex components, *ECT2, ECT4*, and *ECT6*, also decreased following 1 hour of heat stress while showing a notable recovery after 24 hours (Fig. 1C). Interestingly, the eraser enzyme *ALKBH10B* and *ALKBH9B* exhibited increased transcript levels at both 1 and 24 hours of heat stress, suggesting an elevated capacity for m6A removal upon heat stress exposure (Fig. 1C, S Fig. 1). This sharp decrease of m6A immediately after the HS treatment suggests a potential adaptive strategy of the plants to reduce the overall m6A machinery that could influence mRNA stability, translation, or degradation of heat-stress related transcripts.

**Figure 1:**
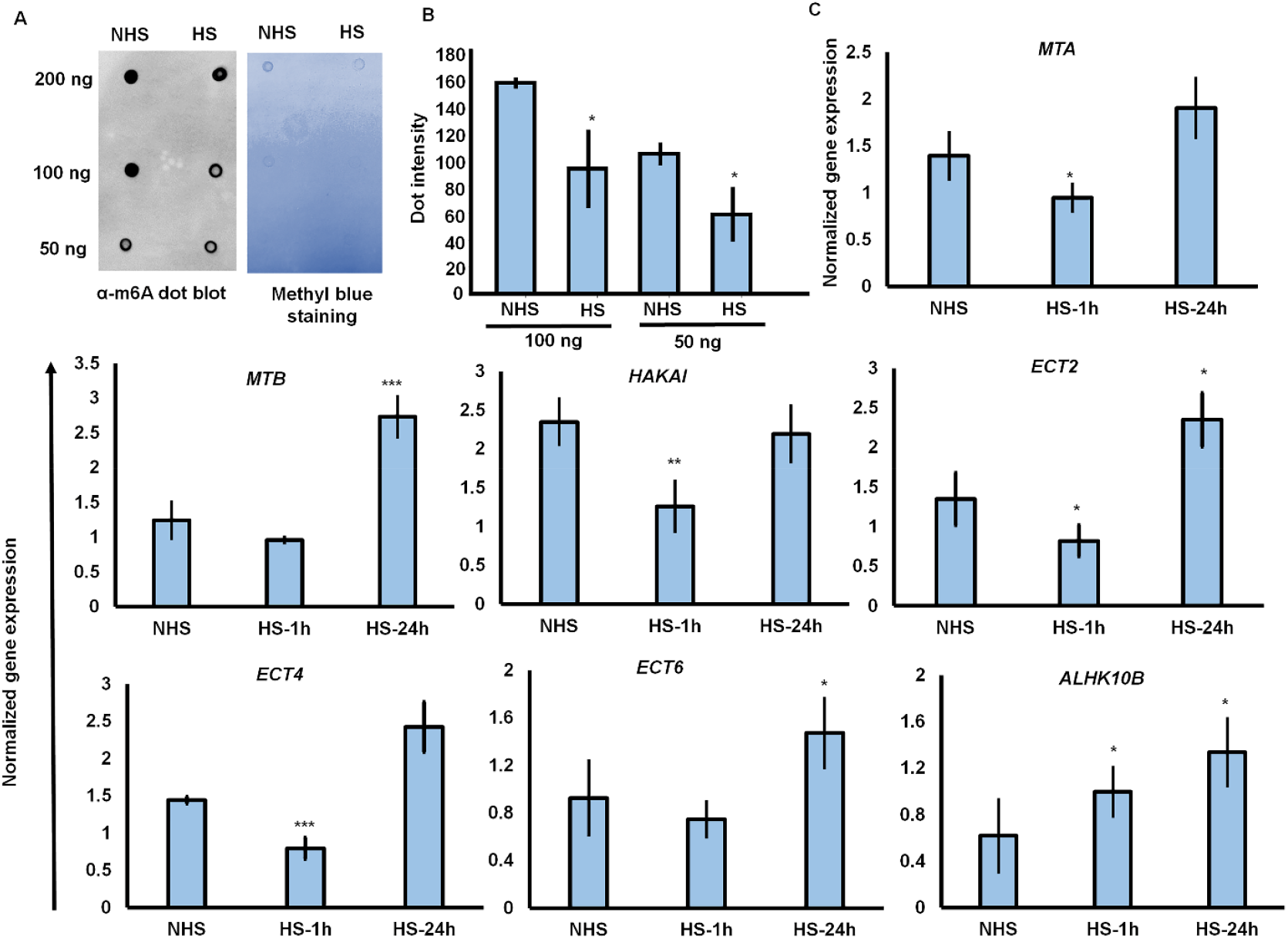
Heat Stress-Induced modulation of m6A machinery. A. Dot blot assay showing the levels of m6A in poly(A+) mRNA isolated from Col-0 seedlings in control (NHS) and after 1 h of HS treatment. Methylene blue staining represents the loading control. B. Quantification of m6A levels by dot intensity. 3 µL dots of 200, 100 and 50 ng poly(A+) from the same replicate are blotted onto the nylon membrane. C. Expression analysis of core Arabidopsis m6A machinery genes in Col-0 seedlings after 1 and 24 hours of heat stress (HS) treatment. This includes m6A writer genes (*MTA, MTB* and *HAKAI*), reader genes (*ECT2,4* and *ECT6*), and eraser genes (*ALKH10B*). Plots represent the mean of 3 biological replicates. Error bars represent SD. Asterisks indicate a statistical difference based Student’s t-test (*P ≤ 0.05; **P ≤ 0.01; ***P ≤ 0.001).

### m6A modification acts as a negative regulator of thermotolerance

To understand the direct role of m6A in the regulation of plant thermotolerance, we took advantage of Arabidopsis lines known to be deficient in m6A methylation such as *mta* carrying a T-DNA insertion in the major methyltransferase. To rescue *mta* from its lethal embryonic developmental defect, *mta ABI3:MTA* was used, expressing *MTA* under the embryo-specific *ABI3* promoter (Bodi *et al*., 2012). In addition, we used an *mta* complementation line *pMTA:MTA-YFP/mta* as control. We exposed 11 days old seedlings to a heat stress of 44°C for 30 minutes. Compared to wildtype and the *pMTA:MTA-YFP/mta* complementation line, the *mta* mutant plant showed significantly less HS induced plant damage as the fresh weight was 72% and 73% higher compared to Col-0 and the *pMTA:MTA-YFP* complementation line, respectively (Fig. 2A-C). Similarly, the *mta* line showed 76% higher survival compared to Col-0 plants (Fig. 2C). The total chlorophyll and ion leakage data further support these findings, showing a 50% increase in total chlorophyll content in *mta* plants compared to wildtype Col-0 (Fig. 2D). Additionally, the *mta* line exhibited significantly lower ion leakage, with a 56% reduction compared to Col-0 (Fig. 2E), indicating improved membrane stability under heat stress conditions. Other m6A lines such as the writer deficient mutant line *fip37-4* (SALK_018636) (Růžička *et al*., 2017) and eraser overexpressor *35S:ALKBH10B* (Prall *et al*., 2023) also showed significantly less damage under HS conditions as compared to Col-0 (Fig. 2F). Moreover, m6A-reader mutants like *ect2* and *ect2,4* double mutant lines also exhibited enhanced thermotolerance compared to wild-type plants (S Fig. 2A, B). Overall, these results reveal a role for m6A RNA methylation in response to plant heat stress resistance.

**Figure 2:**
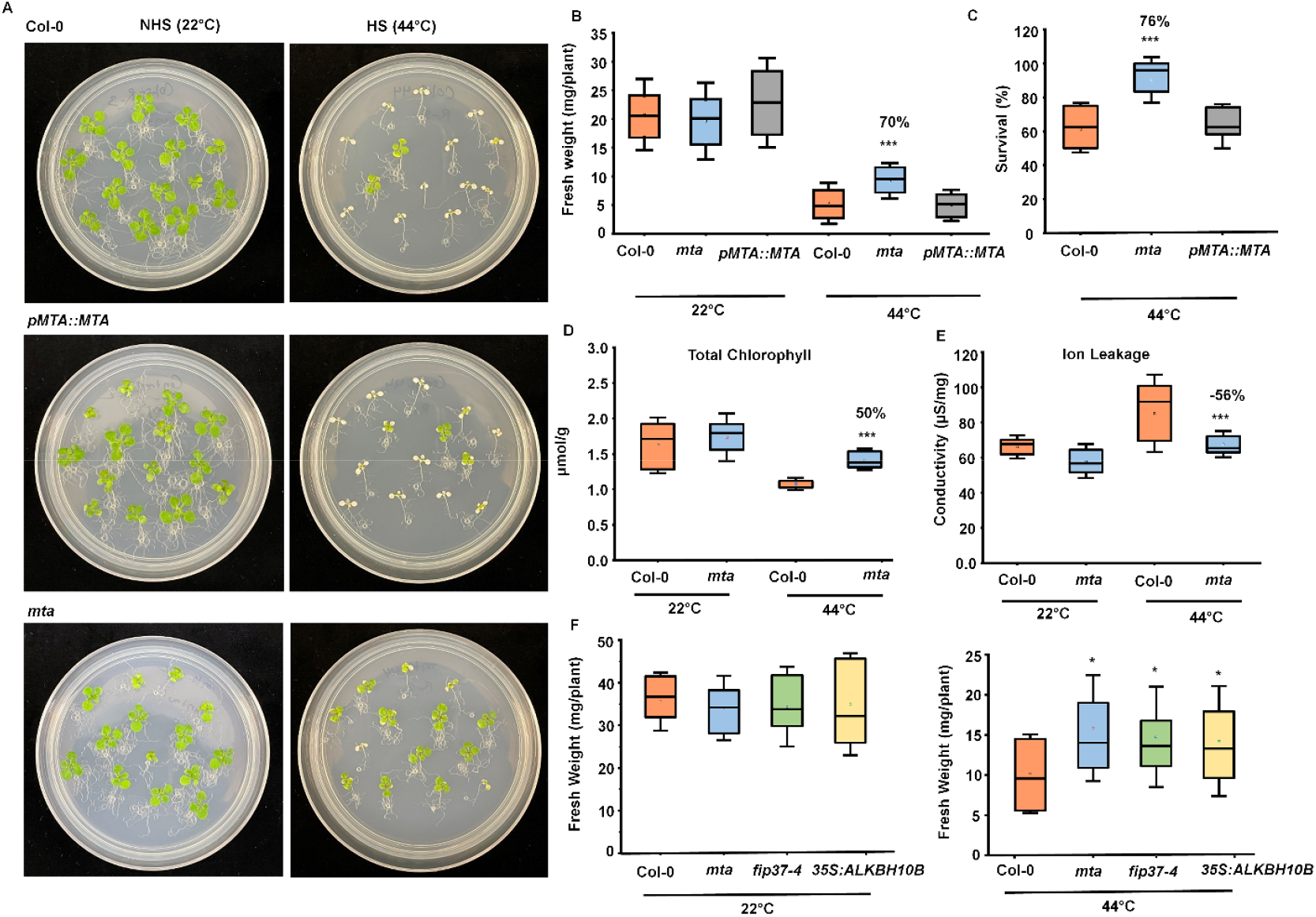
m6A Modification acts as a Negative Regulator of Thermotolerance. A. Phenotypic comparison of wildtype Col-0, *mta* complementation line (*pMTA::MTA*), and *mta* mutant plants under normal (NHS) and heat stress (HS) conditions at 44°C for 30 minutes. B, C. Quantification of fresh weight and percent survival in 11-day-old seedlings exposed to 44°C heat stress (HS) and 22°C normal (NHS) conditions, followed by a 4-day recovery at 22°C. D. Total chlorophyll content in Col-0 and *mta* plants after 4 days of heat stress. E. Ion leakage assessment in Col-0 and *mta* mutant plants under normal (NHS) conditions and after 24 hours of heat stress. F. Plant biomass measurement in Col-0, *mta, fip37-4*, and *35:ALKBH10B* overexpressor lines under normal (NHS) and heat stress conditions, with recovery after 4 days. Plots represent the mean of 3 biological replicates (n = 72, 24 plants/biological repeat. Error bars represent SD. Asterisks indicate a statistical difference based Student’s t-test (*P ≤ 0.05; **P ≤ 0.01; ***P ≤ 0.001).

### *mta* exhibits enhanced transcript levels of heat stress associated genes

Since the *mta* line showed resistance to heat stress, we wanted to compare the basal transcript levels of heat stress signature genes of the *mta* line with those of wild type Col-0. RNA sequencing (RNA-seq) analysis under non-heat stress conditions revealed that the *mta* plants exhibited significantly higher transcript levels of key heat stress-responsive genes including heat stress transcription factors, such as *HSFA2,3, HSP18*.*2, HSP90*.*1, HSP20, HSP70, HSP70b, BIP3* and *ATERDJ3A* (Fig. 3A) (Datasheet 1). These results suggest that m6A modification regulates the expression of key heat stress transcription factors and heat stress protein encoding genes. The higher basal expression levels in *mta* suggest that m6A deficient plants are primed to heat stress, due to their inherent upregulation of genes typically associated with heat stress resilience. We next investigated the expression patterns of 11 heat stress-responsive genes, including *HSFA2, 3, 9, HSP18*.*2, HSP90*.*1, HSP20, HTT3, ERF53, 54, HSP70b, BIP3*. To assess their response to heat stress (HS), we performed qRT-PCR analysis at two time points: at 1 and 24 hours after exposing plants to 44°C for 30 minutes. Under ambient non heat stress (NHS) conditions, the *mta* plants exhibited enhanced transcript levels for all 11 heat-responsive genes compared to Col-0 wild-type plants, confirming the RNA-seq data (Fig. 3B). Moreover, the expression levels of these genes were significantly higher in *mta* plants compared to the wild type after 1 and 24 hours of HS. The elevated transcript levels of the heat-responsive genes in *mta* correlate with the improved survival of these plants under severe HS recovery conditions, suggesting that the observed heat stress phenotype is regulated by the m6A machinery.

**Figure 3:**
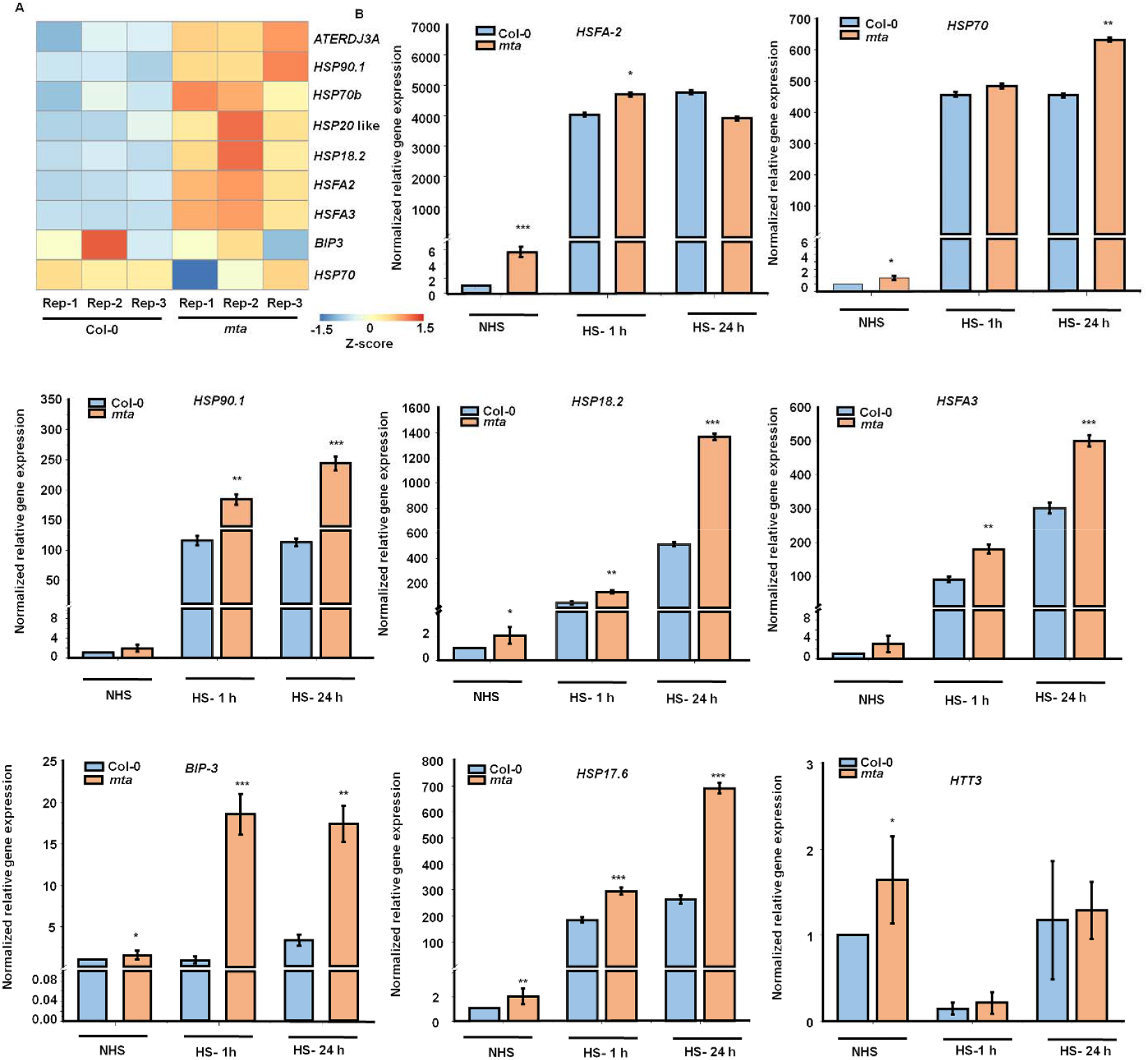
Absence of m6A modification enhances transcript levels of heat stress response genes. A. RNA-seq analysis showing transcript levels of heat stress-associated genes in wildtype Col-0 and *mta* mutant plants. B. Transcript levels of heat stress response genes, including *HSFA2, HSP70, HSP90*.*1, HSP18*.*2, HSFA3, BIP3, HSP17*.*6*, and *HTT3*, in Col-0 and mta mutant plants under control (NHS), after 1 hour, and after 24 hours of heat stress (HS). Plots represent the mean of 3 biological replicates. Error bars represent SE. Asterisks indicate a statistical difference based Student’s t-test (*P ≤ 0.05; **P ≤ 0.01; ***P ≤ 0.001).

### MTA regulates stability of heat stress-associated gene transcripts

Given that m6A modification is known to regulate mRNA stability (Lee *et al*., 2020), we next investigated the influence of m6A modification on the stability of transcripts involved in the heat stress response. After allowing *HSP mRNA* accumulation for 24 hours post heat shock, we inhibited transcription with Actinomycin D and monitored transcript decay for selected transcripts at 0, 3 and 6 hours post-treatment (S Fig. 3). Our results revealed a significantly faster decay of the heat stress-responsive transcripts *HSFA2, HSFA3, HSP101, HSP90*.*1, HSP18*.*2, HSP70, HSP70b, HSP17*.*6*, and *ERF53* in wild-type compared to *mta* mutant plants (Fig. 4A) (S Fig. 3). This accelerated decrease in transcript levels in Col-0 plants suggests that m6A modification enhance the degradation of these mRNAs upon heat stress, thereby contributing to the dynamic regulation of heat stress-responsive genes. In contrast, the slower decay rates observed in the *mta* mutant plants indicate that the absence of m6A modifications stabilize these transcripts leading to their prolonged life span during heat stress. These findings support the idea that MTA regulated RNA decay is a crucial mechanism in fine-tuning gene expression during the heat stress response.

**Figure 4:**
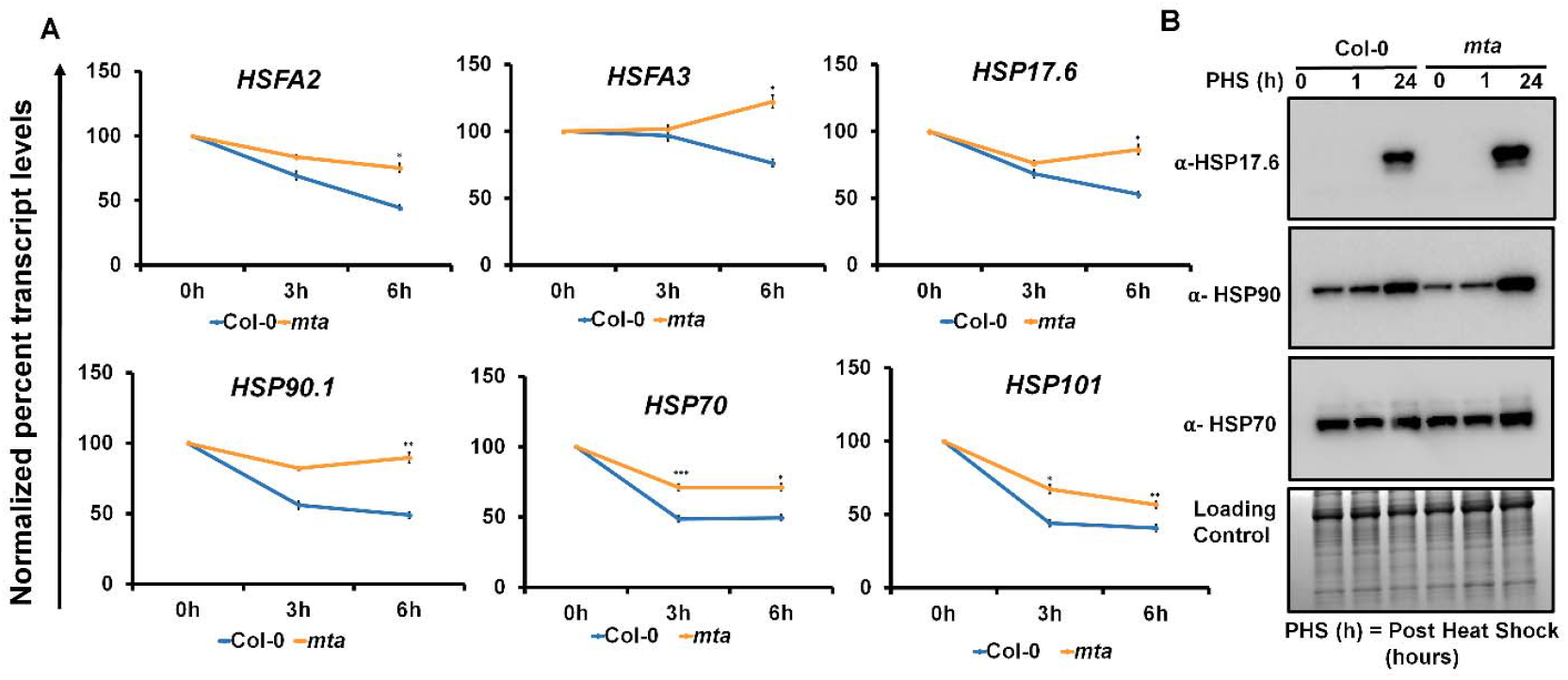
m6A regulates the abundance of heat stress associated gene and protein level, shaping how cells react to temperature changes. A. Actinomycin D–mediated mRNA decay assay: 11-day-old seedlings were transferred to ½ MS liquid medium and subjected to heat stress at 44⍰ °C for 30 minutes. After a 24-hour recovery period, Actinomycin D (30 ⍰ μg/mL; Sigma) was added to inhibit transcription and samples were harvested at 0 (immediately after Act-D addition), 3, and 6 hours post-treatment. The relative transcript levels of HSFA2, HSFA3, HSP17.6, HSP90.1, HSP70, and HSP101 were quantified via RT-qPCR. mRNA levels at time 0 were set to 100% as baseline, and transcript abundance at 3 h and 6 h was calculated relative to this 100% 0 h baseline to assess mRNA decay. B. Western blot analysis: Protein levels of HSP17.6, HSP90, and HSP70 in Col-0 seedlings were analyzed at 0, 1, and 24 hours of heat stress. Ponceau S staining served as a loading control. Error bars represent standard deviation (SD). Asterisks denote statistically significant differences (Student’s t-test: P ≤ 0.05; P ≤ 0.01; *P ≤ 0.001).

To investigate the effect of enhanced accumulation of *HSP* mRNA transcripts due to improved stability and transcription priming in *mta*, we quantified the protein levels of selected heat shock proteins (HSPs) using Western blot analysis. Specifically, we examined the levels of HSP17.6, HSP90.1, and HSP70, which are key proteins involved in the plant heat stress response. Our results revealed that *mta* mutant lines exhibited significantly higher levels of HSP17.6, HSP90.1, and HSP70 compared to wild-type plants (Fig. 4 B). These findings further provide strong evidence that the *mta* mutants not only show enhanced transcriptional activation of heat-responsive genes but also translate this into increased production of key HSPs, which likely contribute to their improved thermotolerance.

### *mta* shows H3K4me3 enrichment, amplifying heat stress resilience

The prominently higher basal transcript levels of key heat stress associated genes in *mta* compared to wild-type plants (Fig. 3) led us to hypothesize that m6A methylation may influence chromatin modifications at heat stress-associated gene loci. This is supported by previous findings that *HSFA2* and *HSFA3* promote the recruitment of H3K4me3, a histone mark linked to active transcription at heat stress memory gene loci (Friedrich *et al*., 2021; Lämke, Brzezinka and Bäurle, 2016; Pratx *et al*., 2023). To test this, we analyzed global H3K4me3 levels via Western blot in *mta* and Col-0 plants. Under non-stress conditions, *mta* showed elevated H3K4me3 levels compared to Col-0 (Fig. 5A,B). Using ChIP-qRT-PCR, we further examined the levels of H3K4me3 at the promoters and gene bodies of key heat stress-responsive genes, including *BIP3, HSP70, HSFA2, HSP17*.*1, HSP18*.*2*, and *HSP101* (S Fig. 4). As compared to Col-0 plants, a marked increase in H3K4me3 deposition was found at these gene loci in the *mta* mutant under non-stress conditions (Fig. 5C).

**Figure 5:**
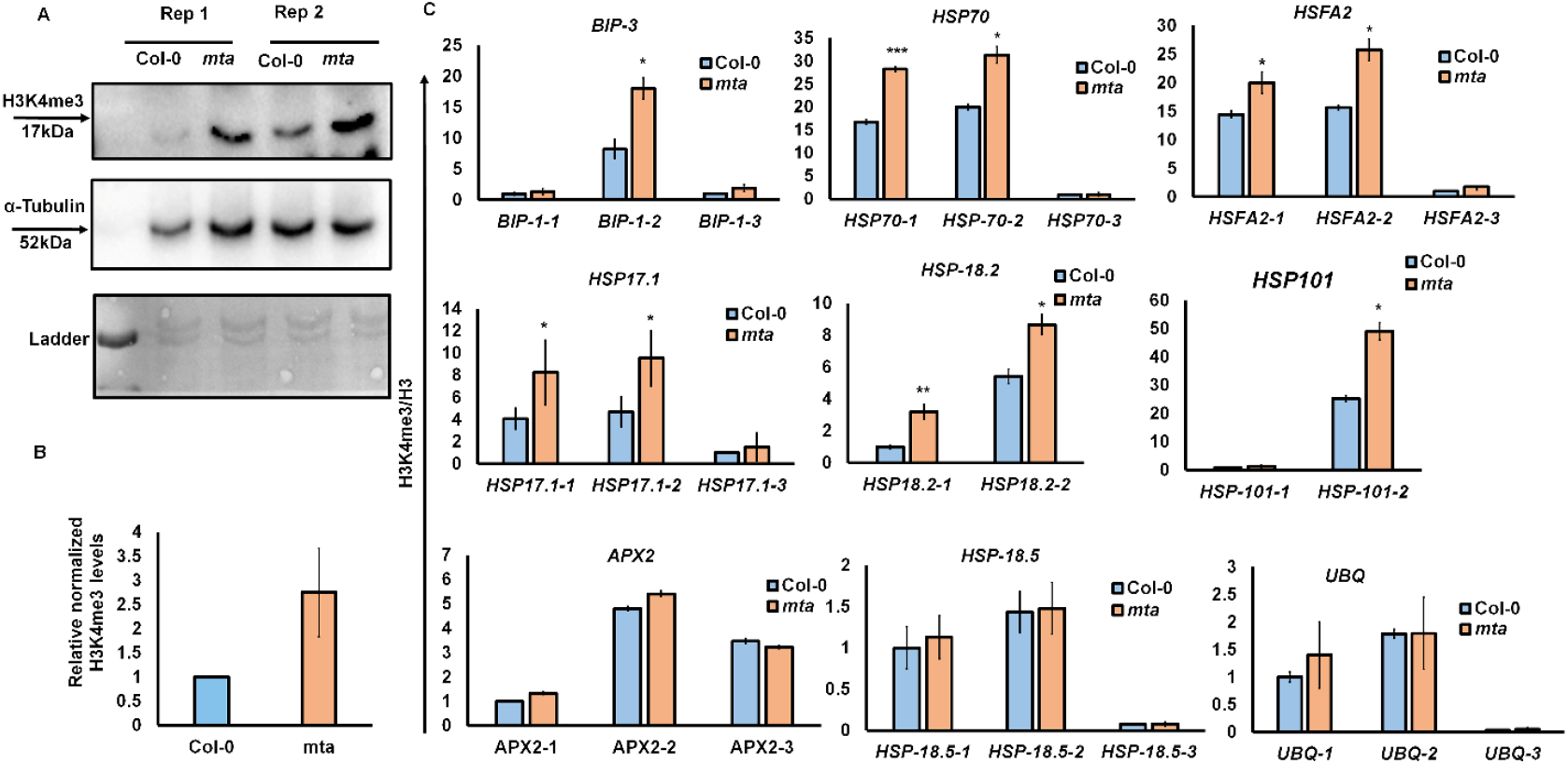
Loss of m6A Modification Leads to Elevated H3K4me3 Levels at Heat Stress-Responsive Genes. A, B. Analysis of global H3K4me3 levels in Col-0 and *mta* mutant plants under normal growth conditions. \C. Enrichment of H3K4me3 marks at heat stress-responsive gene loci in wildtype Col-0 and *mta* plants under control condition (NHS). The genes *APX2, HSP18*.*5*, and *UBQ* were included as negative controls due to their unchanged expression in transcriptomic data, serving to confirm the specificity of H3K4me3 enrichment at heat-responsive genes. Plots represent the mean of 3 biological repeat. Error bars represent SD. Asterisks indicate a statistical difference based Student’s t-test (*P ≤ 0.05; **P ≤ 0.01; ***P ≤ 0.001).

The absence of m6A modification stabilizes the mRNAs of heat stress-responsive genes while also priming their expression through H3K4me3-driven epigenetic changes. The elevated H3K4me3 levels likely facilitate enhanced basal transcription, enabling a more rapid and robust response to heat shock and aiding in the subsequent recovery. This dual role of m6A, influencing both RNA stability and promoting an active chromatin state, indicates its pivotal function in driving robust thermotolerance in plants. We propose an m6A-dependent, multi-layered mechanism that enhances plant resilience to prolonged and severe heat stress, ensuring survival in fluctuating environmental conditions (Fig. 6).

**Figure 6:**
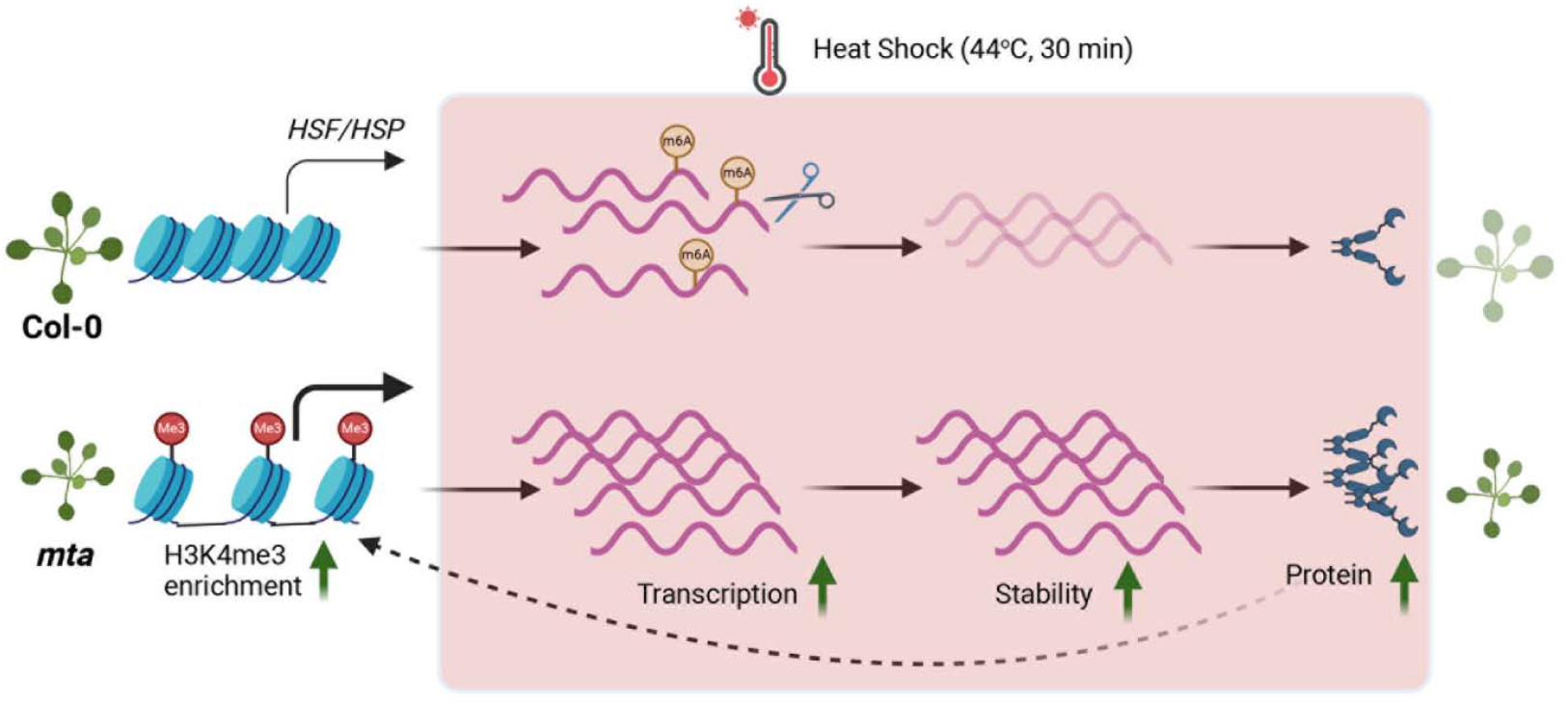
Model of m6A-mediated thermotolerance in Arabidopsis. Under normal, non-heat stress (NHS) conditions, the absence of m6A modification leads to elevated basal expression of heat stress transcription factors, such as HSFA2, in mta plants. This higher expression enhances H3K4me3 enrichment at *HSP* and *HSF* loci, reinforcing a positive feedback loop that primes the expression of *HSPs* and *HSFs*. Under heat shock conditions, this primed state, combined with the increased stability of *HSP/HSF* transcripts in *mta* plants, leads to a substantial rise in HSP protein levels, resulting in enhanced thermotolerance. This model demonstrates that the absence of m6A modification not only stabilizes heat-responsive transcripts but also promotes epigenetic changes that collectively enhance the plant’s ability to withstand high-temperature stress.

## Discussion

N6-methyladenosine (m6A) is a prevalent modification found on most eukaryotic mRNAs. m6A modification is regulated primarily by writer, eraser, and reader proteins and has been shown to play critical roles in various biological processes, including responses to abiotic stresses such as salt, drought, and dark-induced senescence (Anderson *et al*., 2018; Ganguly *et al*., 2024; Hou *et al*., 2022; Hu *et al*., 2021; Jiang *et al*., 2021; Sheikh *et al*., 2024; Zhu, Huo and Pei, 2020). In this study, we explored the role of m6A modification in plant thermotolerance, a significant factor that limits plant productivity and yield. Plants employ several strategies to cope with heat stress. Under extreme high temperatures, plants activate HSFs and produce HSPs, which are essential for their survival (Ohama *et al*., 2017; Ruan *et al*., 2024). Our findings show that m6A machinery mutants exhibit better thermotolerance compared to wild-type plants, as evidenced by their higher survival rates and increased fresh weight after heat stress exposure. This observation aligns with previous research in Drosophila brain cells, where Mettl3 mutant cells (the mammalian homolog of MTA) showed increased resilience to heat stress (Perlegos *et al*., 2022). However, we observed a decrease in global m6A levels at 1 hour post-heat shock, which contrasts with findings in Drosophila and Arabidopsis, where m6A levels increased following heat stress and high light stress (Perlegos *et al*., 2022; Zhang *et al*., 2022). This discrepancy may be due to differences in m6A responses in specific tissues at different time points, such as brain cells in *Drosophila* (Perlegos *et al*., 2022) or differential plant response to intensity and types of stress (Zhang *et al*., 2022). In plants, the observed dip in global m6A levels at 1 hour of HS appears to be coordinated by a decrease in the transcription of writer and reader machinery, while an increase in eraser levels (Fig. 1). Additionally, our results indicate that m6A-mediated thermotolerance in plants may be partly facilitated by reader proteins, as mutants lacking reader proteins like ECT2 and ECT4 also showed increased resistance to heat shock (Fig. 2). This aligns with previous studies in mammals, where reader proteins have been implicated in mediating heat stress responses (Perlegos *et al*., 2022; Zhou *et al*., 2015), and is further supported by findings on plant ECT proteins in regulating mRNA stability (Cai *et al*., 2024; Song *et al*., 2023).

Furthermore, our data demonstrate that *mta* plants have elevated basal mRNA levels of *HSFs* and *HSPs* relative to wild-type plants under normal (NHS) conditions. These mRNA levels not only increase more significantly in response to heat stress but also remain elevated for a longer period in *mta* mutant plants. These effects could arise from m6A-dependent mechanisms, either through pre-transcriptional chromatin modifications or post-transcriptional mRNA processing (Sharma *et al*., 2023). One of the well-documented roles of m6A is to regulate the stability of stress-responsive transcripts, helping to fine-tune gene expression under stress conditions (Arribas-Hernández *et al*., 2018). Using Actinomycin D chase assay, we could show that m6A modification contributes to plant thermotolerance by destabilizing mRNAs. The slower mRNA decay rates of heat-responsive genes in *mta* mutants align with earlier reports that m6A modification promotes RNA degradation via the recruitment of specific RNA-binding proteins, such as YTHDF2 in animals (Wang *et al*., 2014). We could also show that m6A-mediated RNA decay is a critical mechanism in regulating the heat stress response by ensuring timely degradation of heat stress-responsive transcripts. This regulation contributes to a dynamic and fine-tuned gene expression landscape during stress adaptation. Similar mechanism in Drosophila, *mettl3* mutant brain cells, which are deficient in m6A, show increased resistance to thermal and oxidative stress due to the stabilization of stress-responsive transcripts. Similarly, we also observed that *mta* exhibited elevated levels of transcription factors and chaperone proteins, suggesting that the m6A-mediated regulation of gene expression in plants is an essential mechanism to fine-tune the timing and intensity of the heat stress response. By limiting excessive transcription and mRNA turnover, m6A helps to balance the need for the immediate production of stress response proteins. In the absence of m6A, as observed in the *mta* mutant, this regulatory balance is altered, leading to higher basal expression and enhanced thermotolerance due to the increased stability of stress-related transcripts.

In addition to influencing stability, m6A is also known to regulate the translation of heat stress-responsive mRNAs. For instance, in mammalian cell lines, m6A modulates cap-independent translation of HSP70 to enhance the heat stress response (Zhou *et al*., 2015). In our study, we similarly observed increased protein levels of key HSPs following heat stress (Fig. 4C). In this regard, we believe the m6A-mediated transcript stability is a major factor driving the higher protein levels. However, the exact m6A-mediated translation control of plant HSPs needs further investigation.

The observation that *mta* has elevated basal levels of *HSF* and *HSP* transcripts under non stress conditions suggested a role of m6A at the epigenetic level. We could indeed find enhanced H3K4me3 levels in *mta* plants at the gene loci of *HSFA2* and several *HSPs*. This epigenetic process might be facilitated by HSFA2 and HSFA3, which have been shown to promote H3K4me3 enrichment on heat stress memory genes (Friedrich *et al*., 2021; Lämke, Brzezinka and Bäurle, 2016; Pratx *et al*., 2023). Particularly, heteromeric complexes of HSFA2 and HSFA3 drive transcriptional memory in Arabidopsis, enhancing the plant’s ability to withstand future heat stress episodes (Friedrich *et al*., 2021). These modifications facilitate the maintenance of gene expression memory, ensuring a rapid response to subsequent heat stress events. It would be interesting to investigate whether *mta* plants also rely on the HSFA2-HSFA3 axis-driven thermo-priming in future studies. Nonetheless our data suggests a regulatory feedback loop model in which the elevated basal transcript levels of *HSFA2* and *HSFA3* could initiate the enrichment of the H3K4me3 histone marks at the *HSF* and *HSP* gene loci. This histone modification subsequently facilitates a more accessible chromatin state, which further amplifies the heat stress response at the transcriptional level. As a result, the enhanced expression of these transcription factors leads to the sustained induction of *HSPs*, such as *HSP70* and *HSP101*. This cascade effectively reinforces a thermomemory state, ensuring a robust heat stress response and thermotolerance in plants (Fig 6). This study opens up several avenues for future research. For instance, since our analysis focused on seedlings, the combined effects of m6A on epigenetic and transcriptional regulation of thermotolerance may vary across different tissues and developmental stages (Wang *et al*., 2022). Investigating the spatio-temporal impact of m6A on plant thermotolerance could provide further insights into these mechanisms. Additionally, stress granule (SG) formation, a known response to heat stress in plants, is an area where m6A might play a role. As SGs are enriched with m6A-modified mRNAs (Fu and Zhuang, 2020), it would be intriguing to explore whether m6A also regulates SG-mediated thermotolerance, potentially offering new perspectives on how plants manage heat stress at the RNA level (Fu and Zhuang, 2020).

## Data Availability

The RNA-seq data has been submitted to NCBI under Project ID PRJNA1025919.

### Acknowledgments

We thank all the members of the Hirt Lab and the CDA management and greenhouse facility team of KAUST for technical assistance. Special thanks to Naganand Rayapuram for his valuable scientific insights and discussions throughout the work.

## Author contributions

The study was conceived and designed by K.S., A.H.S., and H.H. K.S. standardized the heat stress protocol, phenotyping experiments, performed RT-PCR experiments and analyzed the data. A.H.S. Performed RNA-seq. K.N. performed bioinformatics analysis on the raw RNA-seq; A.P. J helped in bioinformatics analysis. A.F. and W.A. assisted in experiments. K.S. wrote the manuscript. H.H. and A.S. edited the manuscript. All authors approved the final version of the manuscript.

